# Circulating adiponectin mediates the association between omentin gene polymorphism and cardiometabolic health in Asian Indians

**DOI:** 10.1101/2020.08.20.258897

**Authors:** Karani S. Vimaleswaran, Juanjie Jiang, Dhanasekaran Bodhini, Kandaswamy Ramya, Deepa Mohan, Coimbatore Subramanian Shanthi Rani, Nagarajan Lakshmipriya, Vasudevan Sudha, Rajendra Pradeepa, Ranjit Mohan Anjana, Viswanathan Mohan, Venkatesan Radha

## Abstract

**Background:** Plasma omentin levels have been shown to be associated with circulating adiponectin concentrations and cardiometabolic disease-related outcomes. However, the findings have been inconsistent due to high level of confounding. In this study, we aim to examine the association of omentin gene polymorphism with plasma adiponectin levels and cardiometabolic health status using a genetic approach, which is less prone to confounding, and investigate whether these associations are modified by lifestyle factors such as diet and physical activity.

**Methods:** The study included 945 normal glucose tolerant and 941 unrelated individuals with type 2 diabetes randomly selected from the Chennai Urban Rural Epidemiology Study (CURES), in southern India. Study participants were further classified into cardiometabolically healthy and unhealthy, where cardiometabolically healthy were those without hypertension, diabetes, and dyslipidemia. Fasting serum adiponectin levels were measured by radioimmunoassay. The A326T (rs2274907) single nucleotide polymorphism (SNP) in the exon 4 of the omentin gene was screened by polymerase chain reaction-restriction fragment length polymorphism and direct sequencing.

**Results:** The ‘A’ allele of the omentin SNP was significantly associated with lower adiponectin concentrations after adjusting for age, sex, body mass index (BMI), waist circumference (WC) and cardiometabolic health status (p=1.90 x 10^-47^). There was also a significant association between circulating adiponectin concentrations and cardiometabolic health status after adjusting for age, sex, BMI, WC and Omentin SNP (p=7.47×10^-10^). However, after adjusting for age, sex, BMI, WC and adiponectin levels, the association of ‘A’ allele with cardiometabolic health status disappeared (p=0.79) suggesting that adiponectin serves as a mediator of the association between omentin SNP and cardiometabolic health status. There were no significant interactions between the SNP and dietary factors on adiponectin levels and cardiometabolic health status (p>0.25, for all comparisons).

**Conclusions:** Our findings show that adiponectin might function as a mechanistic link between omentin SNP and increased risk of cardiometabolic diseases independent of common and central obesity in this Asian Indian population. Further studies are required to confirm the molecular mechanisms involved in this triangular relationship between omentin gene, adiponectin and cardiometabolic diseases.

## Introduction

The prevalence of cardiometabolic diseases is rapidly increasing in Asian Indians leading to increased morbidity and mortality due to type 2 diabetes (T2D), dyslipidemia and hypertension [1, 2]. Despite lower body mass index (BMI), Asian Indians are characterised by increased plasma insulin levels, insulin resistance, increased waist circumference, excess visceral fat, lower high-density lipoprotein cholesterol (HDL-c), increased triglyceride levels and higher proportion of small dense low-density lipoprotein cholesterol (LDL-c) [2, 3]. In addition, Asian Indians have low adiponectin levels [3], an adipokine with insulin-sensitizing, anti-apoptotic, and anti-inflammatory properties, which has shown to play an important role in the pathogenesis of dyslipidemia by affecting HDL-c and LDL-c metabolism [4–6].

Omentin, also referred as intelectin-1, is another adipocytokine that is highly expressed in human visceral fat tissue [7]. *In vitro* and animal studies have shown that omentin enhances insulin action in human adipocytes and has beneficial effects on cardiovascular system [8, 9]. Majority of the studies have shown that circulating levels of omentin are decreased in individuals with obesity and T2D [10–13] and positively associated with flow-mediated vasodilatation [14, 15] suggesting a protective role of human omentin on cardiometabolic health. Even though omentin and adiponectin have been shown to have anti-inflammatory and cardioprotective effects, only a few studies have examined the physiological link between these adipokines in relation to cardiometabolic diseases [16, 17].

Several studies have demonstrated an association of omentin gene single nucleotide polymorphism (SNP), A326T (rs2274907), in the exon 4 with cardiometabolic disease-related traits. A study in a South Asian population (N=350) [18] has shown a significant association between the SNP rs2274907 and coronary artery disease. In addition, a study in 168 Iranians showed an association of this SNP with BMI and T2D [19]. Furthermore, a study in 495 Central-Europeans showed that the T allele of the SNP rs2274907 was associated with lowest average energy intake (7877 ± 2780 J/day) [20]. However, a few small studies have failed to show a significant association between the SNP and cardiometabolic traits [21, 22], which could be either due to insufficient statistical power or differences in genetic heterogeneity. To date, there are no genetic studies to establish the molecular link between omentin, adiponectin and cardiometabolic health status. Given that genetic associations are less prone to confounding [23, 24], in the present study, we have used an extensively studied missense polymorphism, rs2274907, in the omentin gene, as a genetic instrument to test for its association with serum adiponectin levels and cardiometabolic health status in up to 1,886 individuals from an Asian Indian population. In addition, we have tested for the interaction of this genetic variant with lifestyle factors such as diet and physical activity on adiponectin levels and cardiometabolic health.

## Materials and Methods

### Study population

The study participants were chosen from the urban component of the Chennai Urban Rural Epidemiology Study (CURES), a cross-sectional epidemiological study conducted on a representative sample of the population of Chennai in Southern India [25]. The details of the study has been published elsewhere [25]. Briefly, in phase 1, 26,001 individuals were recruited based on a systematic random sampling technique. Participants with selfreported diabetes taking drug treatment for diabetes were classified as “known diabetes”. All individuals with known diabetes (n = 1,529) were invited to visit the center for detailed studies. In addition, every 10th individual of the 26,001 individuals without known diabetes was invited to undergo oral glucose tolerance tests using a 75-g oral glucose load (dissolved in 250 ml of water) (Phase 3 of CURES). Those who were confirmed by oral glucose tolerance test to have 2-h plasma glucose value ≥ 11.1 mmol/l (200 mg/dl) based on World Health Organization (WHO) consulting group criteria were labelled as “newly detected diabetes” and those with 2-h plasma glucose value<7.8 mmol/l (140 mg/dl) as being normal glucose tolerant (NGT) [26]. For the present study, 945 NGT and 941 unrelated individuals with T2D and genetic data were included. Informed consent was obtained from all study participants, and the study was approved by the Madras Diabetes Research Foundation Institutional Ethics Committee.

### Phenotype measurements

Anthropometric measurements including weight, height, and waist circumference (WC) were obtained using standardized techniques. The BMI was calculated as weight (in kg) divided by the square of height (in m). Blood pressure was recorded in the sitting position in the right arm to the nearest 2mmHg using the mercury sphygmomanometer (Diamond Deluxe BP apparatus, Pune, India). Two readings were taken 5 minutes apart and mean of two was taken as the blood pressure. Biochemical analyses were done on a Hitachi-912 Auto Analyzer (Hitachi, Mannheim, Germany) using kits supplied by Roche Diagnostics (Mannheim). Fasting plasma glucose (glucose oxidase–peroxidase method), serum cholesterol (cholesterol oxidase-phenol-4-amino-antipyrene peroxidase method), serum triglycerides (glycerol phosphatase oxidase-phenol-4-amino-antipyrene peroxidase method), and HDL-c (direct method; polyethylene glycol-pretreated enzymes) were measured. LDL-c was calculated using the Friedewald formula [27]. Glycated haemoglobin (HbA1c) was estimated by high-performance liquid chromatography using a Variant™ machine (Bio-Rad, Hercules, CA, USA). Serum insulin concentration was estimated using an enzyme-linked immunosorbent assay (Dako, Glostrup, Denmark).

### Assessment of serum adiponectin concentrations

Fasting adiponectin levels were measured using radioimmunoassay (Cat. No. HADP-61HK, Linco Research, St Charles, MO, USA). The intra-assay and the inter-assay co-efficient of variation were 3.8 and 7.4 per cent respectively and the lower detection limit was 1 ng/ml [28]. Adiponectin data was available for 1,205 samples.

### Dietary intake and physical activity assessments

Dietary intakes were assessed using a previously validated and published [29] interviewer administered semi-quantitative food frequency questionnaire (FFQ) containing 222 food items to estimate food intake over the past year. A detailed description of the development of FFQ and the data on reproducibility and validity had been published [29]. A validated self-report questionnaire was used to measure physical activity [30]. For the present study, only a subset of the study population (N=513) had data on dietary intake and physical activity.

### Definitions

Study participants were classified into cardiometabolically healthy (N=370) and unhealthy (N=1,516), where cardiometabolically healthy were those without hypertension, diabetes, and dyslipidemia [31]. Hypertension was diagnosed in all subjects who were on drug treatment for hypertension or if the blood pressure ≥ 140/90 mmHg [32]. National Cholesterol Education Programme guidelines were used to define those with and without dyslipidemia [33].

### SNP Genotyping

The SNPrs2274907 was genotyped by polymerase chain reaction on a GeneAmp PCR system 9700 thermal cycler (Applied Biosystems, Foster City, CA) using the primers, forward: 5’ CCTCTGCAGATCCAAAGGTG 3’ and reverse: 5’ CCGCACTGAGAATGGTGTTA 3’. The PCR amplicons were digested with AccI restriction enzyme (New England Biolabs, Inc., Beverly, MA) and the resulting products were electrophoresed on a 3% agarose gel. Based on the analysis of 200 blind duplicates (20%), there was 100 % concordance in the genotyping. Furthermore, a few variants were confirmed by direct sequencing with an ABI 310 genetic analyzer (Foster City, CA). The SNP was in Hardy Weinberg equilibrium (HWE) (P=0.59).

### Statistical analyses

Descriptive statistics are presented as means and SD for continuous variables and as percentages for categorical variables. To test whether the observed genotype counts were in HWE, a goodness-of-fit chi-square test was performed. Student t test as appropriate was used to compare groups for continuous variables. Given the low frequency of the rare homozygotes, dominant model was used (comparing individuals with common homozygous genotypes with the combined group of rare homozygotes and heterozygotes). The genetic associations with the continuous and categorical outcomes were examined using linear and logistic regression models, respectively, adjusting for age, sex, BMI and serum adiponectin, wherever appropriate. Interactions between the SNP and dietary intake were assessed by including an interaction term in the linear and logistic regressions. All analyses were carried out using SPSS, version 26. A P value <0.05 was considered to be statistically significant.

### Power calculation

Given that there are no previously reported effect sizes for the associations and interactions pertaining to the Omentin SNP and adiponectin concentrations in the Asian Indian population, we were unable to perform a prospective power calculation. However, based on the most significant associations observed in the present study, we performed a retrospective power calculation using QUANTO software, Version 1.2.4 (May 2009). We performed power calculations in the form of least detectable effects based on the assumption of significance levels and powers of 5 and 80%, respectively. At 80% power, the minimum detectable effect was beta 1.50 μg/mL (adiponectin concentrations) for a SNP with minor allele frequency of 21% in the case-control analysis (N=1,204).

## Results

### Associations between cardiometabolic health status and clinical and biochemical parameters

In the present study, 80.3% of the individuals were cardiometabolically unhealthy with significantly higher BMI, WC, fasting plasma glucose, insulin, total serum cholesterol, LDL-c, triglycerides, systolic and diastolic blood pressures and HbA1c and lower HDL-c and serum adiponectin concentrations (p<9.6 x 10^-9^ for all comparisons) (**Table 1**).

**Table 1:** Clinical and biochemical characteristics of the participants from the CURES study

| **Clinical and biochemical parameters** | **Cardiometabolically healthy (N=370)** | **Cardiometabolically Unhealthy (N=1,516)** | **P value** |
| --- | --- | --- | --- |
| **Age (years)** | 37±13 | 47±13 | 3.41 x 10<sup>-36</sup> |
| **Sex (men/women)** | 111/259 | 736/780 | 1.27 x 10<sup>-10**</sup> |
| **Body mass index (kg/m<sup>2</sup>)** | 22.1±4.8 | 24.9±4.4 | 3.39 x 10<sup>-22</sup> |
| **Waist circumference (cm)** | 79±12 | 89±11 | 1.58 x 10<sup>-41</sup> |
| **Waist-hip ratio** | 0.85±0.09 | 0.92±0.08 | 1.32 x 10<sup>-49</sup> |
| **Fasting plasma glucose (mg/dl)** | 84±8 | 134±67 | 1.30 x 10<sup>-146</sup> |
| **Fasting serum insulin (μIU/ml)** | 7.6±5.2 | 10.1±6.5 | 4.33 x 10<sup>-13</sup> |
| **Systolic blood pressure (mmHg)** | 116±16 | 126±20 | 5.46 x 10<sup>-24</sup> |
| **Diastolic blood pressure (mmHg)** | 73±10 | 77±11 | 2.77 x 10<sup>-9</sup> |
| **Total serum cholesterol (mg/dl)** | 162±20 | 196±43 | 3.20 x 10<sup>-92</sup> |
| **Serum triglycerides (mg/dl)** | 79±25 | 163±109 | 4.68 x 10<sup>-137</sup> |
| **Low-density lipoprotein cholesterol (mg/dl)** | 97±18 | 122±37 | 6.55 x 10<sup>-72</sup> |
| **High-density lipoprotein cholesterol (mg/dl)** | 50±7 | 41±10 | 5.98 x 10<sup>-61</sup> |
| **Glycated hemoglobin (%)** | 5.5±0.4 | 7.6±2.4 | 7.23 x 10<sup>-179</sup> |
| **Fasting serum adiponectin (µg/mL)** | 9.9±5.6 | 7.4±4.3 | 9.60 x 10<sup>-9</sup> |
| **Total energy intake (kcal)** | 2655±579 | 2813±919 | 0.12 |
| **Carbohydrate energy (%)** | 65±7 | 65±6 | 0.97 |
| **Protein energy (%)** | 11±1 | 11±1 | 0.35 |
| **Fat energy (%)** | 23±5 | 23±5 | 0.47 |
| **Fibre (g/day)** | 3±8 | 33±12 | 0.07 |
| **Physical activity levels (Sedentary %/**<br>**Moderately active %/ Physically active %)** | 81/ 18/ 1 | 82/ 15/ 3 | 0.52** |

Data shown are represented as means ± SD, wherever appropriate
\*P values generated from an independent samples 't' test for the differences in the means/proportions between cardiometabolically healthy and unhealthy participants
\*\* P value generated from a chi-square test

There was a significant association between circulating adiponectin concentrations and cardiometabolic health status after adjusting for age, sex, BMI, WC and Omentin SNP rs2274907, where cardiometabolically unhealthy individuals had 2.00 μg/mL decrease in adiponectin concentrations compared to the control group (p=7.47×10^-10^).

### Association between the SNP and serum adiponectin concentrations

The Omentin SNP rs2274907 was significantly associated with serum adiponectin concentrations after adjusting for age, sex, BMI, WC and cardiometabolic health status (p=1.90 x 10^-47^), where ‘A’ allele carriers (AT + AA) had significantly lower levels of adiponectin (means ± SE: 4.2 ± 2.7 μg/mL) compared to those with TT genotype (means ± SE: 8.9 ± 4.5 μg/mL) (**Figure 1**).

**Figure 1:**
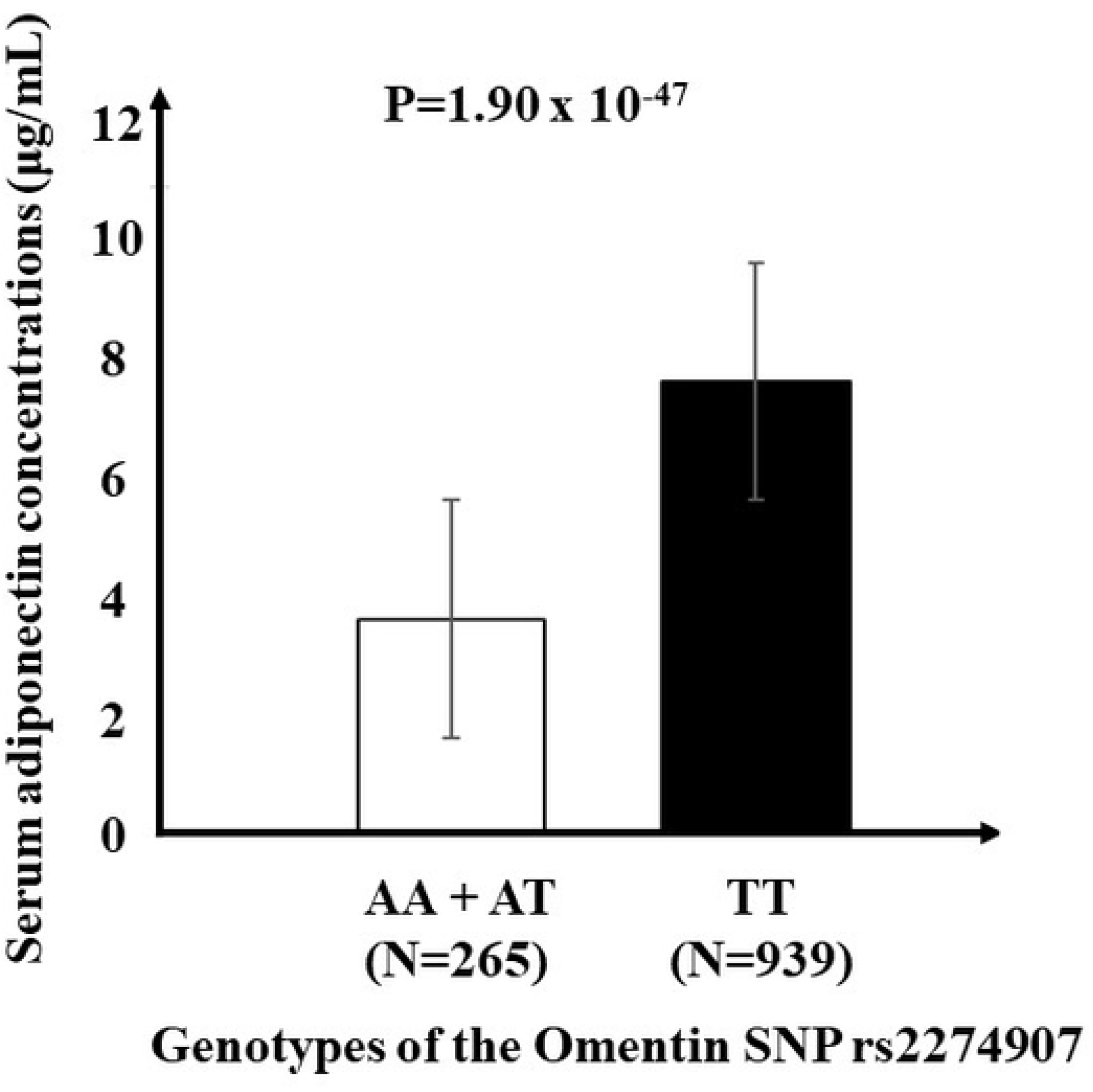
Association between omentin SNP rs2274907 with serum adiponectin concentrations. After adjusting for age, sex, BMI, WC and cardiometabolic health status, the ‘A’ allele carriers (AA + AT) have significantly lower levels of serum adiponectin concentrations compared to those with TT genotype (p=1.90 x 10^-47^). Abbreviations: BMI, Body mass index; WC, waist circumference; SNP, Single nucleotide polymorphism.

### Association between the SNP and cardiometabolic health status

After adjusting for age, sex, BMI and WC, there was a significant association between the ‘A’ allele of the SNP rs2274907 and increased risk of cardiometabolic health status, where ‘A’ allele carriers had 1.35 times increased risk of being cardiometabolically unhealthy compared to TT homozygotes (p=0.03). However, after adjusting for age, sex, BMI, WC and adiponectin concentrations, the ‘A’ allele was not associated with cardiometabolic health status (P=0.79) (**Figure 2**).

**Figure 2:**
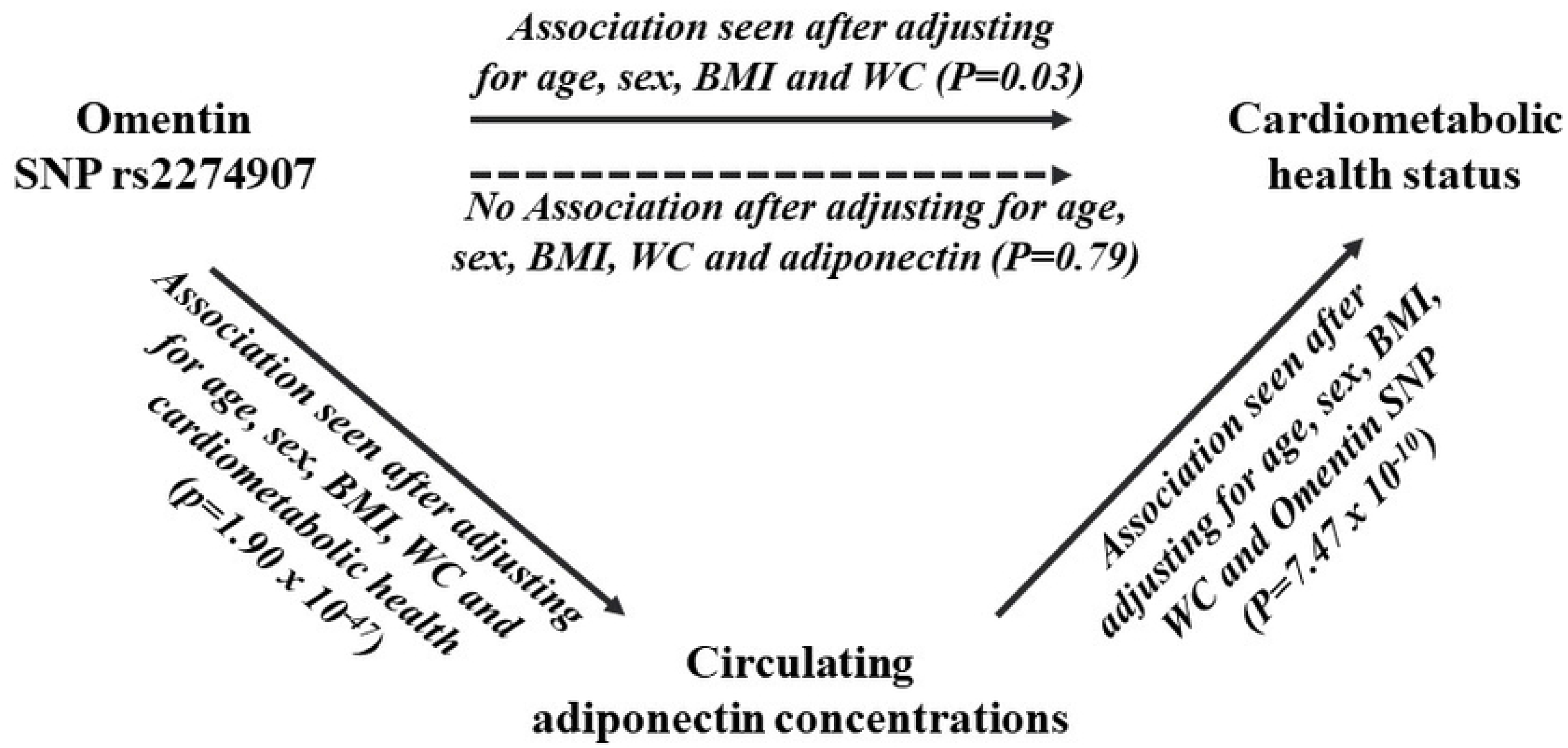
Diagram showing the role of circulating adiponectin as an intermediate predictor between omentin SNP and cardiometabolic health status. One-sided arrows with unbroken lines represent significant associations and one-sided arrows with broken lines represent lack of associations. There is a significant association between the omentin SNP rs2274907 and lower serum adiponectin concentrations after adjusting for age, sex, BMI, WC and cardiometabolic health status (p=1.90 x 10^-47^). There is a significant association between serum adiponectin concentrations and cardiometabolic health status after adjusting for age, sex, BMI, WC and SNP rs2274907 (p=7.47×10^-10^). There is no association between the omentin SNP rs2274907 and cardiometabolic health status after adjusting for age, sex, BMI, WC and serum adiponectin concentrations (p=0.79). Abbreviations: BMI, Body mass index; WC, waist circumference; SNP, Single nucleotide polymorphism.

There was also no significant difference in the genotype and allele frequencies of the SNP between cardiometabolically unhealthy and healthy individuals under an additive (p=0.31) and a dominant (p=0.40) model (**Supplementary table 1**).

### Interaction between SNP and adiponectin concentrations on cardiometabolic health status

To test if the association between the SNP and cardiometabolic health status is modified by serum adiponectin concentrations, we examined the interaction between the SNP and adiponectin on cardiometabolic health status. There was no evidence for a significant interaction (p_interaction_=0.24) suggesting that adiponectin is unlikely to modify the effect of the SNP on cardiometabolic health status.

### Interaction between SNP and lifestyle factors on adiponectin concentrations and cardiometabolic health status

There was no significant interaction between the SNP and lifestyle factors such as carbohydrate energy %, fat energy %, protein energy %, dietary fibre intake and physical activity levels on adiponectin concentrations and cardiometabolic health status (P>0.25, for all comparisons), respectively (**Table 2**).

**Table 2:** Interactions between the Omentin SNP rs2274907 and lifestyle factors on adiponectin concentrations and cardiometabolic health status.

|  |  |  |  |  |
| --- | --- | --- | --- | --- |
| *P values for the interactions between the Omentin SNP and dietary factors on serum adiponectin concentrations\*\** |  |  |  |  |
| SNP*Carbohydrate<br>energy % | SNP*Protein energy<br>% | SNP*Fat energy<br>% | SNP*Fibre<br>(g)<sup>‡</sup> | SNP*physical<br>activity |
| 0.77 | 0.37 | 0.57 | 0.98 | 0.95 |
| *P values for the interactions between the Omentin SNP and dietary factors on cardiometabolic health status\*\*\** |  |  |  |  |
| SNP*Carbohydrate<br>energy % | SNP*Protein energy<br>% | SNP*Fat energy<br>% | SNP*Fibre<br>(g)<sup>‡</sup> | SNP*physical<br>activity |
| 0.87 | 0.26 | 0.14 | 0.30 | 0.69 |

\*\*P values obtained from linear regression analysis after adjusting for age, sex, body mass index, waist circumference and cardiometabolic health status
\*\*\* Data are p values obtained from logistic regression analysis after adjusting for age, sex, body mass index, waist circumference and adiponectin concentrations
<sup>‡</sup> Adjusted for age, sex, body mass index, waist circumference, adiponectin concentrations and total energy intake
SNP, Single Nucleotide Polymorphism

## Discussion

Our study provides the first evidence for the role of circulating adiponectin as a mechanistic link in the association between omentin SNP rs2274907 and cardiometabolic health independent of common and central obesity in this Asian Indian population. In this study, we observed a strong cross-sectional association between the omentin SNP and serum adiponectin concentrations, even after accounting for the potential confounders including cardiometabolic health status. Similarly, a strong association was seen between adiponectin concentrations and cardiometabolic health status after adjusting for the confounders including the omentin SNP. However, the association between ‘A’ allele of the omentin SNP rs2274907 and cardiometabolic health disappeared after adjusting for serum adiponectin concentrations, which suggests that the association is likely to be mediated through circulating adiponectin. Hence, modulating adiponectin concentrations through lifestyle interventions might be an effective approach to overcome the genetic risk of cardiometabolic diseases.

Adiponectin, an adipokine with 244 amino acids, is synthesized by adipocytes and has been shown to stimulate the glucose uptake in skeletal muscles and decrease the hepatic glucose synthesis [34], whereas omentin, a 313-amino-acid-long polypeptide adipokine, is mainly synthesized by visceral adipose tissue and is involved in cellular energy homoeostasis and vascular tone regulation [8]. Both animal and human studies have demonstrated a positive correlation between omentin and adiponectin concentrations [12, 35–37]. However, there is no study, to date, that has examined the genetic variants in omentin gene with serum adiponectin concentrations. The present study has examined the A326T (SNP rs2274907), which is a single nucleotide missense polymorphism in the exon-4 of the Omentin 1 gene, which substitutes valine instead of aspartic acid at position 109 (Val109Asp or V109D). Even though the position 109 is located outside the fibrinogen domain of Omentin protein, several studies have reported association of the SNP rs2274907 with cardiometabolic diseases [18–22, 38] suggesting that the SNP might be in linkage disequilibrium (LD) with another causative variant. Interestingly, a study in 4,200 North Indians that had examined 207 common genetic variants in proximal promoter and untranslated regions of genes on 1q21-23 and 20q13 identified omentin SNP rs1333062 as one of the top signals for T2D [38]; this SNP in the 3’ flank region of omentin gene is in complete LD with Val109Asp - SNP rs2274907 (D’=1.0 and r^2^=1.0 for Gujarati Indians in Houston, Texas population from HapMap data). Although the amino acid position 109 does not represent a completely conserved consensus position [5], the direct adjacent site at 108 contains a highly conserved amino acid (alanine) and hence, the proximity of the substituted amino acid at 109 position to the highly conserved amino acid at 108 might be of functional relevance.

Our study has shown that the minor ‘A’ allele of the omentin SNP rs2274907 is associated with cardiometabolic health under the mediation of circulating adiponectin levels. The effect of the minor allele on chronic disease outcomes has also been demonstrated in other ethnic groups including Pakistani (N=350) [18], Iranian (N=168 & 282) [22, 39] and Turkish (N=87) [40] populations. However, a couple of small studies, one in a Caucasian population (N=390) and one in an Indian population (N=500) [41], failed to provide an evidence of an association of the SNP with T2D [21]. These discrepancies are likely to be due to the existence of genetic heterogeneity across different ethnic groups and insufficient statistical power to detect small effect sizes. To date, the present study has been the largest (N=1,886) to explore the association of the omentin SNP rs2274907 with cardiometabolic diseases. While majority of the studies have shown ‘T’ allele as the minor allele [18, 22, 39, 40], the dbSNP database shows ‘A’ allele as the minor allele except for some of the Asian, African and European populations, which is indicative of a significant genetic heterogeneity (https://www.ncbi.nlm.nih.gov/projects/SNP/snp_ref.cgi?do_not_redirect&rs=rs2274907). However, ‘A’ allele is the minor allele in our study and the dbSNP database for GIH population (Gujarati Indians in Houston, Texas, USA). This suggests that future large studies focusing on omentin SNP rs2274907 is highly warranted in each ethnic group to understand the ethnic-specific role of the variant in cardiometabolic diseases.

Several studies have implicated hypoadiponectinemia in the pathogenesis of T2D and cardiovascular diseases [42–44]. Higher concentrations of adiponectin have been shown to decrease the cardiometabolic risk [45, 46], given its role in promoting anti-inflammatory effects, improving insulin sensitivity, increasing glucose uptake by the cells and producing endothelial nitric oxide [47, 48]. While some studies have shown adiponectin as an independent risk factor for cardiometabolic diseases [43, 49], there are a few studies that have shown adiponectin as a mediator of the cross-talk between adipose tissue and cardiovascular system [50]. Furthermore, a few studies have also explained the role of adiponectin as a mediator in the relationship between omentin and cardiometabolic diseases [35, 51]. This finding is in line with our study in Asian Indians, where circulating adiponectin levels serve as a mediator in conferring the genetic risk of cardiometabolic diseases. A previous study in a Spanish population has shown a significant association of the minor allele of the omentin SNP rs12409609, which is in complete LD with SNP rs2274907, with low omentin gene expression [52]. Given the positive correlation between plasma omentin and adiponectin levels [12, 35–37], it is possible that low omentin expression might contribute to the reduction in adiponectin levels, which, in turn, could possibly increase the cardiometabolic risk among ‘A’ allele carriers. However, mechanistic studies are required to confirm the triangular relationship between omentin gene, adiponectin and cardiometabolic risk.

Our study findings are suggestive of the fact that modulation of adiponectin levels might be an effective strategy to overcome the genetic risk of cardiometabolic diseases. Previous studies have examined the impact of lifestyle modifications and drug therapies to improve the circulating adiponectin concentrations [53–55]. A few studies have shown that weight loss [56] and combined diet control and physical exercise [57] can increase plasma levels of adiponectin, while smoking [58] and increasing activity of the sympathetic nervous system [59] can decrease the adiponectin concentrations. Drugs such as renin-angiotensin system blocking agents [60], peroxisome proliferator activated receptor (PPAR)-alpha agonists [61], PPAR-gamma agonists [62] and hypoglycemic drugs [63] have been shown to improve adiponectin concentrations. To understand whether modifying the lifestyle could overcome the genetic risk of hypoadiponectinemia and cardiometabolic diseases, we investigated the SNP-lifestyle interactions and found that none of the interactions were statistically significant. The lack of interaction could be a result of the small sample size and insufficient statistical power to detect the small effect sizes of the interactions.

The main strength of the study is the large sample size from a well characterised population, which is representative of the city of Chennai, and the study is sufficiently powered to detect the genetic associations. The other strength is the use of a validated FFQ, which has shown high reproducibility and validity for total carbohydrates and dietary fibre. However, our study has a few limitations which need to be acknowledged. We performed a cross-sectional study and hence, we are unable to infer causality between the SNP, adiponectin concentrations and cardiometabolic disease outcomes. Although confounders were adjusted in our regression analyses, we cannot exclude the residual confounding due to unknown factors. Another limitation is the recall bias from FFQ which cannot be ruled out. Even though our study is sufficiently powered to detect the genetic associations, the sample size is small for detecting significant gene-lifestyle interactions, which might be the reason for the lack of significant interactions. Furthermore, omentin protein or mRNA expression levels were not assessed in the study and hence, it is not possible to confirm whether the SNP has any influence on the omentin gene expression. Finally, given that the outcome is a multifactorial trait, the present study has examined only one genetic variant from the omentin gene; however, this is the only coding region variant that has been extensively studied in the gene.

In summary, we have identified a robust association between the omentin SNP and serum adiponectin concentrations and the latter with cardiometabolic disease outcomes, suggesting that adiponectin could be a pathogenic mediator of the genetic susceptibility towards cardiometabolic disease outcomes. These findings suggest that targeting adiponectin might be beneficial in overcoming the genetic risk of cardiometabolic diseases. Hence, lifestyle interventions and drug therapies to increase adiponectin levels could serve as effective tools in preventing cardiometabolic diseases. However, mechanistic studies are required to confirm this epidemiological relationship before strategies to promote adiponectin modulation could be implemented.

## Supplementary Materials

**Supplementary table 1:** Association between the Omentin SNP rs2274907 and cardiometabolic health status

## Acknowledgements

We thank all study participants for their cooperation. Dr. Karani S Vimaleswaran acknowledges support from the British Nutrition Foundation (BNF). This is the 157^th^ paper from CURES (CURES-157).

## Authors’ contributions

VKS conceived the study, performed the statistical analysis and drafted the manuscript; JJ and BD assisted with the statistical analysis and writing of the manuscript; MV, ARM, DM, SCS, PR and SV designed the CURES study; VKS designed the nutrigenetics study; RV designed the genetic study; RK performed the genotyping analysis; PR, ARM, DM, LN, SV, SK, MV and RV critically reviewed the manuscript. All authors contributed to and approved the final version of the manuscript.

